# Measuring the Immune Memory Response of In Vitro Polarized Th1, Th2, and Th17 Cells in the Face of OVA Transgenic *Leishmania major in* Mouse Model

**DOI:** 10.1101/2024.07.10.601391

**Authors:** Mebrahtu G. Tedla, Musammat F. Nahar, Alison L. Every, Jean-Pierre Y. Scheerlinck

## Abstract

Th1 and Th2 cytokines determine the outcome of *Leishmania major* infection and immune protection depends mainly on memory T cells induced during vaccination. This largely hinges on the nature and type of memory T cells produced. In this study, transgenic *Leishmania major* expressing membrane associated ovalbumin (mOVA) and soluble ovalbumin (sOVA) are used as a model to study whether fully differentiated Th1/ Th2 &Th17 cells can recall immune memory and tolerate pathogen manipulation. Naïve OT-II T cells were *in vitro* polarised into Th1/Th2, and these cells were transferred adoptively into recipient mice. Following transferring the memory cells, recipient mice were challenged with OVA transgenic *Leishmania major* and wild type parasite was used a control. The *in vitro* polarised T helper cells continued to produce the same cytokine signatures after challenged by both forms of OVA-expressing *Leishmania major* parasites *in vivo*. This suggests antigen-experienced cells cells remain the same or unaltered in the face of OVA transgenic *Leishmania major*. Such ability of the antigen-experienced cells to remain resilient to manipulation by the parasite signifies that vaccines might be able to produce immune memory responses and withstand against the parasite immune manipulation and protect the host from infection.

## 1. Introduction

*Leishmania major* causes a wide spectrum of disease outcomes ranging from asymptomatic to fatal infection. This is not only due to the variations in the pathogenicity of the parasite strains but also due to a difference in the susceptibility of the host (1). Early immune response during *Leishmania* infection determines the progression of the diseases either acute(self-clearing) or chronic. To address these, various experimental studies have been conducted in mouse strains. Following *L. major* infection, some of the mouse strains develop CD4+ Th1 cell-mediated resistance whereas other strains show a CD4+ Th2 mediated response and they are very susceptible to infection (2). For example, amongst mouse strains, C57BL/6 mice can resolve infections and produce long-lasting immunity, while BALB/c mice are unable to control infections leading to disease progression and in most cases. The ability of C57BL/6 mice to control infection is linked to the production of Th1 cytokines (3). The expression of IFN-γ is shown to be associated with resolving infection in human (4,5) and murine *Leishmaniasis* (6). Macrophage activation also plays a crucial role and IFN-γ is required as a macrophage-activating factor during elimination of *Leishmania* (7,8). IL-4 is also important in macrophage activation (9) and in the upregulation of CR3, a macrophage receptor for *Leishmania* (10,11). Acquired resistance to *L. major* infection relies on the activation of CD4^+^ T cells, which leads to the production of large amounts of IFN-γ and this induces parasite killing by macrophages following nitric-oxide release (12–14). Though the killing of the parasite is mainly orchestrated by IFN-γ through direct macrophage activation (6–8), the direct inhibition of the IL-4(15,16) and the reduction in clonal expansion of Th2 cells by IFN-γ (17) is also important. In addition, direct mechanisms allowing IL-4 to inhibit IFN-γ-activity have been identified, including the direct inhibition of macrophage activation to eliminate intracellular amastigotes 18). Although CD4^+^ T cells produce IL-10, which is important in promoting parasite survival, how such responses develop from precursors during infection is not known (19). Therefore, one can conclude that both Th1 and Th2 cytokines contribute in opposing ways to the pathogenic outcome of *Leishmania* infection (20).

Despite the promising progress in developing vaccines against *Leishmania*, a complete immune protection is not easy to achieve due to lack of persistent parasites and sterile immunity. This is due to parasite antigens are not effective in inducing long term immune memory. Another challenge in achieving vaccine mediated immune protection is the lack of immunodominant antigens that can be recognized by CD4+ T cells (2). Immune memory produced after secondary infection with *L. major* is important in conferring and maintaining protection through central memory CD4^+^ T cells. Indeed, these memory T cells become tissue-homing effector cells that can induce protection against infection. Therefore, immune protection against *L. major* could be mediated through memory T cells induced during vaccination and this largely depends on the nature and type of memory T cells produced (21). During pre-existing immunity, secondary site of parasite infection led to IFN-γ & monocyte mediated parasite killing (22). Studies in resistant C57Bl/6 mice showed pre-existing, effector CD4 T cells (CD44^+^CD62L^−^T-bet^+^Ly6C^+^ effector (T_EFF_)) generated by ongoing chronic infection and they are short-lived if there is no infection (23). Indeed, if the immune effector cells are of the Th1 type and they can withstand any manipulation by the parasite, they may induce protection. If at the other hand, Th2 cells are induced or the Th1 immune responses are manipulated by the parasite to produce Th2 cytokine during the recall response, it is possible that disease might be exacerbated. Therefore, it is essential to establish whether the induced immune memory is resilient to manipulation by the parasite. In this study, OVA-transgenic *L. major* parasite expressing OVA as either a membrane-bound or soluble form, was used as a tool to address aspects of CD4^+^ T cell activation, immune memory resilience and pathogen-mediated manipulation in the OT-II TCR-transgenic mouse model.

## 2. Materials and Methods

### 2.1. Experimental mice

Six to eight-week-old female OT-II and congenic CD45.1 mice, which were used in the present study as sources of naive OT-II T cells and for the adoptive transfer of polarized antigen-experienced cells, respectively were purchased from Walter and Eliza Hall Institute of Medical Research, Melbourne, Australia. All mouse-related research projects require approval from the University of Melbourne’s Animal Ethical Committee (Ethics ID: 1814548.1). OT-II mice are OVA-specific, MHC class II restricted αβ TCR transgenic mice. These OT-II transgenic mice express the mouse alpha-chain and beta-chain T cell receptor that pairs with the CD4 co-receptor and is specific for chicken ovalbumin _323-339_ peptide in the context of I-Ab (CD4 co-receptor interaction with MHC class II).

### 2.2. Leishmania major culture

To produce transfected parasites, *Leishmania major* strain MHOM/SU/73/5-ASKH was employed as the wild type and axenic promastigotes were routinely cultivated in SDM-79 media supplemented with 10% FCS at 27 °C. Parasite cells were cultivated to mid-log phase and extracted by centrifugation (2000 x g, 10 min) for long-term storage. At 10^8^ cells/mL, parasites were resuspended in a mixture of 90% FCS and 10% DMSO. Each cryo tube contained about 2 x 10^7^ cells, which were progressively frozen at -80 °C for overnight before being transferred to liquid nitrogen.

### 2.3. Cloning of OVA gene into *L. major* expression vector pIR1SAT

The gene in sOVA that codes for full-length OVA was digested to get a size of 1158 bp using SmaI and XhoI. The Klenow enzyme was used to produce blunt ends. In order to prevent vector re-ligation, the plasmid vector pIR1SAT (24,25), also known as pSAT, was linearized using SmaI and treated with 1 U of shrimp alkaline phosphatase per μg of DNA. Plasmids were obtained from ampicillin-resistant clones and screened by restriction digestion with EcoRV, which produced a 771 bp DNA fragment from insert ligation into vector with the proper orientation. One EcoRV restriction site on the vector and one on the insert’s surface made it possible to screen for ampicillin-resistant clones that were correctly put into the vector. Additionally, transformants were identified by colony PCR utilizing forward and reverse primers created on the vector backbone and seen by agarose gel electrophoresis in ampicillin-resistant clones. To use for *L. major* transfection, pSAT and pSAT-OVA were digested with SwaI and the targeted fragment was extracted by gel extraction (26).

### 2.4 Genetic manipulation of *L. major*

SwaI was used to digest pSAT and pSAT-derived plasmids to extract the desired area for transfection. With a few adjustments, the fragment of the right size that corresponded to the gene of interest and the targeting region for the ssu gene was electroporated into *L. major* using gel purification (27,28). Centrifugation (1000 x g, 10 min) was used to collect mid-log phase parasites (0.8 x 107 cells/mL). After being washed in 20–30 mL of cold electroporation buffer (1000 x g, 10 min), a total of 4 x 107 cells per transfection were resuspended in 500 μL of cold electroporation buffer. Cuvettes were filled with the prepared cell suspension, and two 1.7 kV, 25 F capacitance electrophoretic pulses were administered. *L. major* was transfected with 5 μg of DNA or no DNA (control). The recovery of parasites took place in 10 mL of SDM-79 media supplemented with 100 μg/mL of penicillin/streptomycin for 24 hours at 27 °C. The selection pressure for the pSAT plasmids was then maintained by dilution of the parasite cultures with 15 mL of SDM-79 medium and nourseothricin (Werner BioAgents, Germany, 110 μg/mL).

### 2.5. Confirming OVA expression by *L. major*

Promastigotes were suspended in 200 µL of lysis buffer (Thermo Fisher Scientific) once they reached the log phase, and 2×106 live promastigotes were then heated to 65 oC for 10 minutes. Using light microscopy, the parasite lysate was centrifuged at 9,400x g for 2 minutes to ensure parasite death. When complete breakdown was attained, the parasite homogenates were subjected to ultrasonic-dependent electronic pulses once more utilizing the microson^TM^ ultrasonic cell disruptor. The supernatants were then purified by passing them through a 0.2 µm filter following high-speed centrifugation. To confirm the above-described protein expression, parasite lysates were separated using SDS-PAGE and then submitted to western blotting using anti-OVA antibodies.

### 2.6. Lymphocyte preparation

Maxillary, inguinal, and splenic lymph nodes of OT-II and CD45.1 were combined in RPMI-1640 (Invitrogen, Life Technologies). A single cell suspension was prepared from the collected lymph nodes for an antigenic stimulation of the whole fraction or non-purified cells. For all the assays, CD45.1 and CD45.2 cells were stained after counting their proportion by flow cytometry from the whole extracted cells. Forceps were used to crush the tissues, and a 70 µm nylon cell strainer was used for the filtering process. The cells were centrifuged at 276xg, for six minutes. After aspirating the supernatants, distilled water was used to lyse the red blood cell pellets for 9 seconds, after which 10% of 10x PBS was added to stop the lysis. Cell suspension was centrifuged at 276 x g, for six minutes. The cells were resuspended in new complete RPMI-1640 media supplemented with 2 mM L-glutamine, 100 U/mL penicillin, 100 g/mL streptomycin, 10% v/v heat-inactivated FCS, and 50 μM 2-Mercaptoethanol after the supernatants had been removed. Cell concentrations were calculated where necessary using a hemocytometer.

### 2.7. *In vitro* polarization of OT-II T cells

OT-II cells were polarized in vitro as previously described (29). The following differentiation cytokines were used to polarize naive OT-II cells into three lineages: 2×10^5^ cells per well were cultured for 4 days and polarized into Th1 (using 10μg/ml OVA peptide, 10 ng/mL of IL-2), Th2 (using 10μg OVA peptide and 10 ng/mL of IL-2, 10 ng/mL of IL-4 and 10 μg/ml of anti-IFN-γ, and Th17 (using 10 μg/ml OVA peptide, 10 μg/ml of anti-IL-4, 10 μg/ml of anti-IFN-γ antibodies, 5 ng/mL of anti-TGF β and 20 ng/mL of IL-6). Naive cells were activated with 10 μg/ml of OVA peptide. To create memory cells, cells were further stimulated for two days with 4.5 ng/mL of IL-7 (Pepro Tech, Rocky Hill, NJ, USA). Th-polarized cells were stimulated with OVA peptide (10 μg/mL) for 72 hours using 10^5^ cells per well. Using flow cytometry, the level of polarization was assessed based on the cytokine profiles of the differentiated cells.

### 2.8. Adoptive transfer of OT-II T cells to congenic mice and parasite injection

According to the manufacturer’s instructions, cells were stained with carboxyfluorescein-diacetate-succinimidyl-ester (CFSE; Bio Legend, San Diego, California, USA). 2×10^6^ CD45.2-expressing OT-II T cells were adoptively transplanted into CD45.1 congenic mice. As a route of infection, we used the subcutaneous (sc) method of administration under the tail base of the recipient mice. Each animal received a s.c. injection of 5×10^6^ empty vector control transformed promastigotes (EVC) and OVA-expressing *Leishmania* promastigotes. As a negative control, cells labeled with CFSE but not challenged with OVA (20 ng/mL) were employed. As a negative control, unchallenged OT-II T cells were employed. Different antigens were administered to mice after 24 hours. Mice were killed after 7 days in order to collect OT-II T cells from the spleen for analysis.

### 2.9. T cell proliferation assay

Cells were extracted from the spleen, maxillary, axillary, and inguinal lymph nodes of OT-II mice using complete RPMI-1640 media (30). The tissue was run through a 70 µm nylon cell strainer to create single cell suspensions. RBCs were lysed according to the previously mentioned method. Following this, cells were labeled with CFSE in accordance with the manufacturer’s instructions. A total of 2×10^5^ CFSE-labeled cells per well were co-cultured for 4-5 days with OVA transgenic or wild type log phase parasite promastigotes. OT-II cells were collected and washed with FACS buffer (10% FCS in PBS) after four days of culture. Following an Fc γR-surface block (anti-Fc γR, clone 2.4G2-16, WEHI Monoclonal Facility), cells were stained with PE-anti-mouse CD4 (Bio Legend) for analyzing proliferation. Prior to data collection with a BD FACSVerseTM flow cytometer (BD Bioscience, New Jersey, USA), 7-amino-actinomycin D (1 µg/ml; Sigma Aldrich, Steinheim, Germany) was added to the cell suspension in order to remove dead cells. Similar to this, adoptively transferred antigen-experienced cells were collected from recipient mice’s spleens after a week, and their proliferation was examined following an ex vivo parasite restimulation.

### 2.10. Cytokine assay

A mouse Th1, Th2, and Th17 cytometric bead array (BD^TM^) was used to measure the cytokine production in cell culture supernatants from T cell proliferation tests. The detection limits for each cytokine were as follows: IL-2 (0.1pg/ml), IL-4 (0.03pg/ml), IL-6 (1.4pg/ml), IFN-γ (0.5pg/ml), TNF (0.9pg/ml), IL-17A (0.8pg/ml), and IL-10 (16.8pg/ml), according to the manufacturer.

### 2.11. Statistical Analysis

The single cell analysis program Flowjo version 10 (Ashland, Oregon, USA) was used to compute and analyze the data. One-way ANOVA was used for all statistical analyses using GraphPad Prism version 7.00 for Windows (GraphPad Software, La Jolla, California, USA). The confidence level was maintained at 95% for all analyses, and a significant threshold of p ≤ 0.05 was needed.

## 3. Results

### 3.1. Cloning of sOVA and mOVA in *Leishmania* expression vector(pIR1SAT)

Since transfection with the targeting region of the plasmid permits recombination of the desired gene into the parasite chromosome and stable expression, the *Leishmania* expression vector, pIR1SAT, was used to express OVA (SERPINB14). As a result, the sOVA.CI plasmid was digested using the restriction enzymes XhoI and SmaI to produce a 1158 bp fragment that matched OVA and contained the N-terminal signal sequence for secretion. This fragment was subsequently gel-purified using a gel purification kit from Promega in accordance with the manufacturer’s instructions (Figure 1A & B). The Klenow enzyme treatment blunted the ends of the purified sOVA fragment. To construct the vector, pSAT-sOVA, the *Leishmania* expression vector pIR1SAT, also known as pSAT, was digested with SmaI and the gene encoding the sOVA expressing fragment gene was introduced using conventional ligation procedures. The ligated DNA was utilized to transform *E. coli* TOP10, and colony PCR was used to check for clones having the right insert. In order to find clones with the insert in the right orientation, plasmid DNA was extracted from positive clones and screened using EcoRV digestion (Figure 1C & D). The sequence encoding the N-terminal 139 amino acids of sOVA were replaced with the sequence encoding the *Leishmania* transferrin receptor in the construct responsible for membrane localization, which was used to express membrane-associated OVA (mOVA).

**Figure 1.**
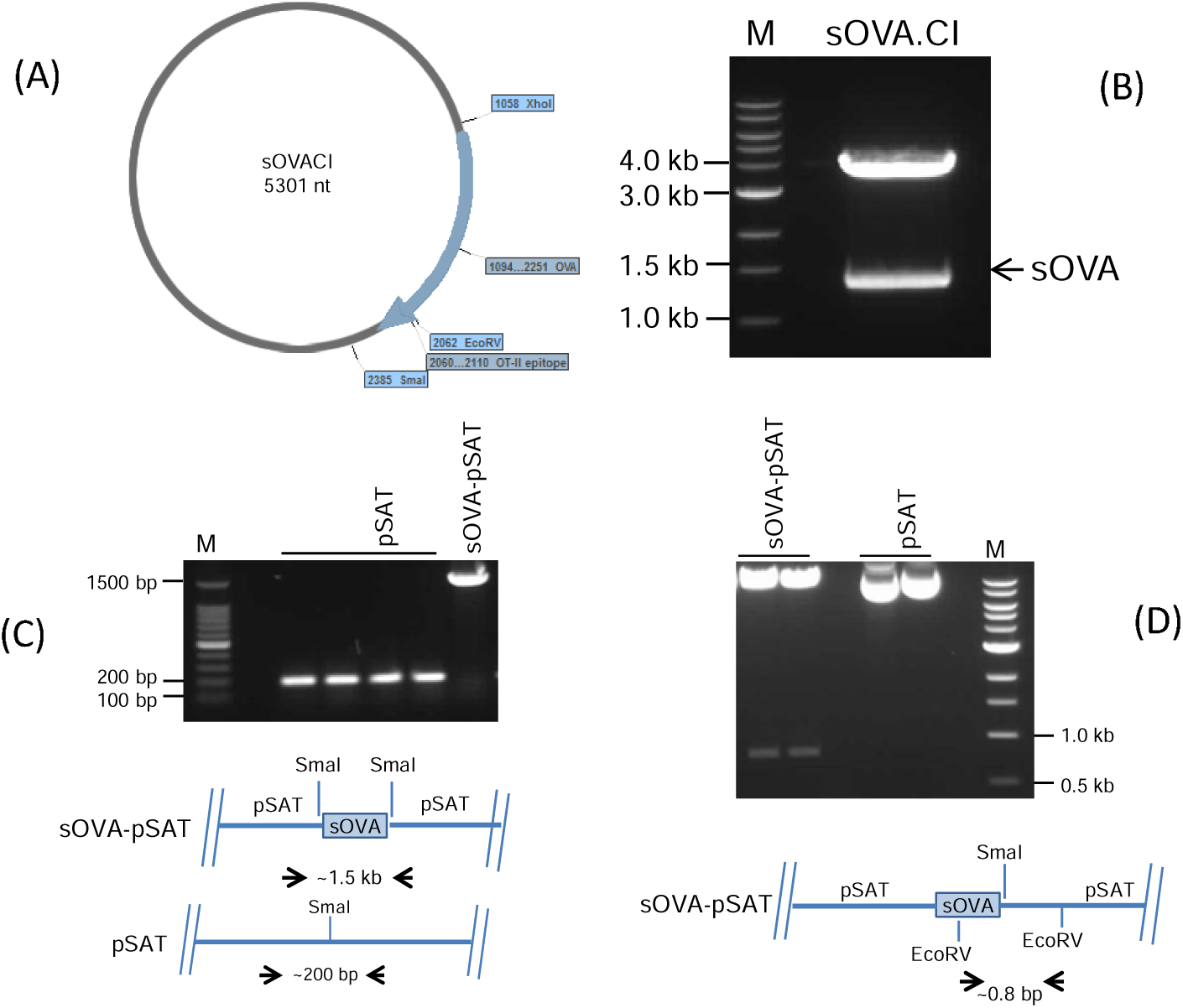
Restriction digestion of sOVA.CI plasmid and cloning of sOVA fragment into pSAT vector. (A) Restrictive digestion was used to remove the sOVA fragment from sOVA.CI plasmid. (B) sOVA.CI was digested by XhoI and SmaI.Cl. Arrow denotes a fragment that was later gel-purified and corresponds to the DNA responsible for encoding sOVA. sOVA, OVA that has been secreted; M, molecular weight marker; sOVA.CI, plasmid carrying sOVA. sOVA fragment cloning into pSAT vector. (C) Following colony PCR using forward and reverse primers created on the pSAT vector backbone, gel electrophoresis shows amplification of the sOVA-pSAT positive clone (2 kb). (D) Due to the presence of one EcoRV site on the sOVA insert and one on the pSAT vector, restriction digestion of the recombinant sOVA-pSAT plasmid with EcoRV produced a 0.8 kb fragment of the positive clones with correct orientation. M, molecular weight marker; pSAT , only pSAT vector; sOVA, secreted OVA; and sOVA-pSAT, sOVA in pSAT vector.

### 3.2. Expression of OVA by transfected *L. major*

To create parasites (and control lines) that stably produce the OVA protein, the *L. major* parasites were transfected with the pSAT-mOVA, pSAT-sOVA, and pSAT plasmids. In order to achieve this, these plasmids were digested with SwaI to produce fragments comprising the 5’ and 3’ targeting areas (homologous to 18s rRNA SSU) and containing the DNA encoding mOVA, sOVA, and pSAT plasmid solely, respectively. These pieces were electroporated into the parasite chromosome and then replaced one 18s rRNA SSU fragment. By using PCR amplification, it was confirmed that the recombinant DNA had been successfully transfected into the parasite (Figure 2A). The vector backbone was amplified by PCR using only the pSAT transfection, which resulted in a 200 bp fragment, as opposed to the expected 1.5 kb and 1.3 kb pieces from the sOVA- and mOVA-containing pSAT constructs. PCR amplification of the inserted DNA using primers flanking the region of insertion, confirming integration into the right location of the genome, further established the OVA-expressing parasite created by the integration of the gene into the parasite chromosome (Figure 2B). The expected 2 kb and 1.8 kb fragments for pSAT-sOVA and pSAT-mOVA, respectively, were produced by the PCR process. *L. major* pSAT (only pSAT DNA transfected), *L. major* pSAT-sOVA (pSAT-sOVA encoding DNA transfected), and *L. major* pSAT-mOVA (pSAT-mOVA expressing DNA transfected) were the three parasite strains that were produced.

**Figure 2.**
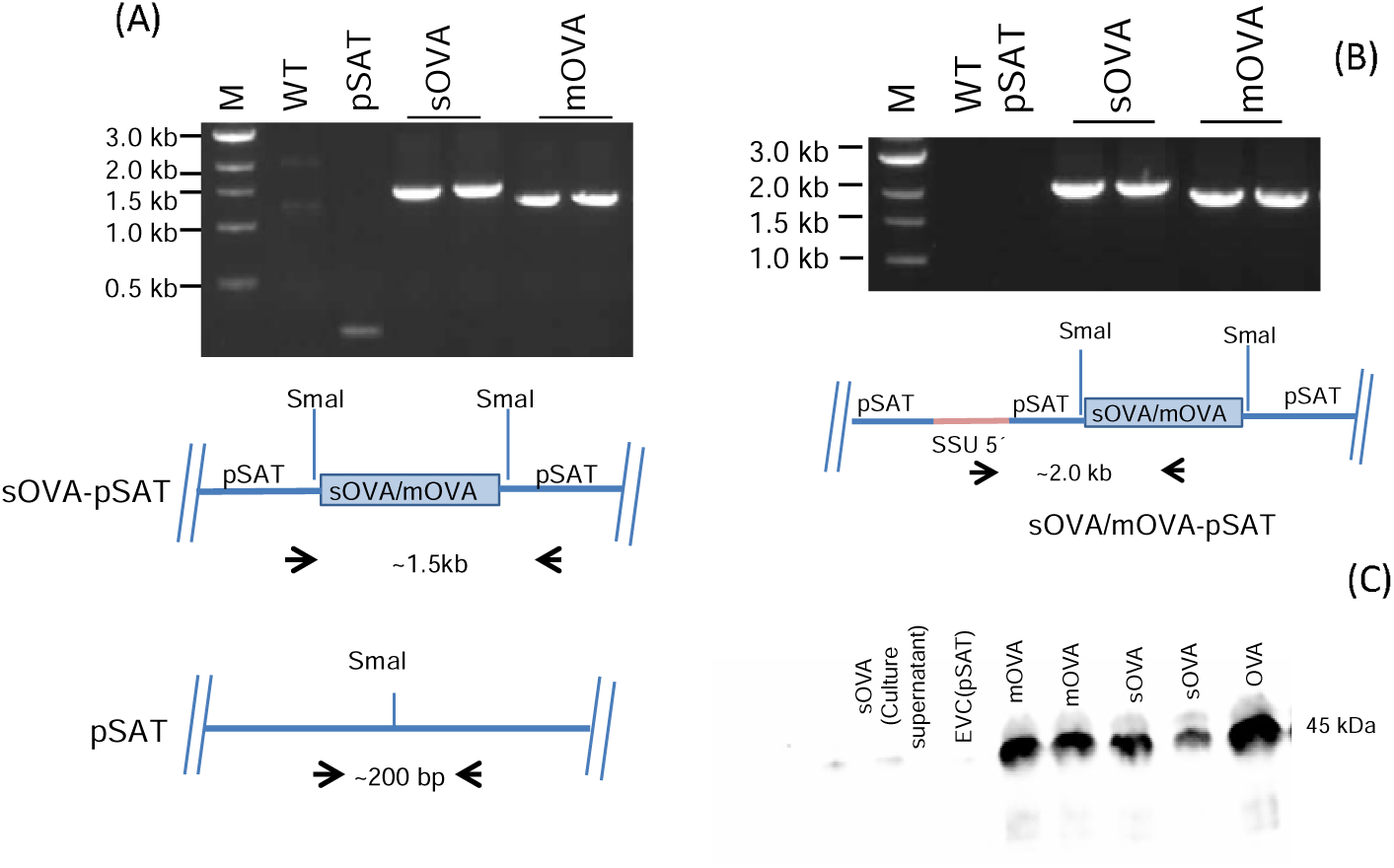
Transfection of *L. major* with sOVA- and mOVA-encoding DNA sequence containing recombinant pSAT. (A) OVA was detected in the genomic DNA that was taken from the transfected *L. major*. Both forward and reverse primers were created using the pSAT vector backbone to conduct the PCR experiment. (B) Using primers based on the *Leishmanial* SSU region (forward) and the middle of the OVA fragment (reverse), the integration of the OVA transgene into the *Leishmanial* chromosome was verified. M stands for molecular weight marker, WT for wild type, pSAT for pSAT vector only, SSU 5’ for small subunit of the *Leishmanial* chromosome 18s rRNA, sOVA-pSAT for sOVA in pSAT vector, and mOVA-pSAT for mOVA in pSAT vector. (C) Confirmation of OVA expression; whole parasite lysates were confirmed using SDS-PAGE followed by subjecting to anti-OVA immunoblotting. A total of 2×10^7^ log-phase promastigotes were lysed to detect protein expression. Both membrane-associated OVA and soluble OVA are detected in the transformed parasites. OVA protein and parasite empty vector control was used as positive and negative control respectively. The first two lanes contain culture supernatants of log-phase *L. major* promastigotes.

Transfection of *L. major* parasites resulted in the successful expression of two forms of OVA protein. The first form of OVA is secreted OVA (sOVA) and the second form of OVA is a membrane-associated OVA (mOVA), which is membrane anchored via a hydrophobic transmembrane sequence. Following transfection of the mid-log phase promastigotes with pSAT-mOVA, pSAT-sOVA and empty vector control or pSAT plasmids, the successful expression of OVA protein was confirmed using western blot (Figure 2C). To confirm the expression of OVA protein using western blot, a pool of approximately 2×10^6^ promastigotes were cultured until they reach into the mid-log phase and a total of 2×10^7^ parasites were lysed using the heat shock method (56 °C for 10 minutes) and the protein was subjected to SDS-PAGE, transferred to a nitrocellulose membrane and detected using OVA-specific antibodies. The supernatant was also assayed separately, but no band was detected (Figure 2C). Promastigotes transfected with pSAT-mOVA and pSAT-sOVA expressed the protein having a band size of 45kda (Figure 2C), which is the same size as the chicken OVA protein. The empty vector control here, referred to as pSAT-plasmid, was used as a negative control and showed no expression of OVA by the parasite.

### 3.3. OT-II CD4^+^ T cells Proliferation by OVA-expressing *L. major*

The *in vitro* activation of OT-II CD4^+^ T cells after exposing the cells to OVA-expressing parasites was determined following extraction of cells from spleen and lymph nodes of OT-II mouse. The results shown in Figure 3, demonstrate that cells stimulated with mOVA expressing parasites had a higher proliferation level (21.8%) than those exposed to sOVA-expressing parasites (18.7%). Cells exposed to OVA peptide (10 μg/ml) and OVA protein (10 μg/ml) showed 61.3% and 44.4% proliferation respectively. To ascertain whether the presence of *L. major* affects OT-II T cell proliferation, OVA peptide and OVA protein was mixed with empty vector control and the response compared to the proliferation in the absence of lysate. The results show that the level of proliferation 61.5% vs 61,3% for OVA peptide and 44.2 % vs 44.4%, for OVA protein was not affected by the presence of the parasite (Figure 3). Some marginal background proliferation was revealed when cells were exposed to parasites transfected with empty vector controls and cells were left unstimulated (3.9 % and 2.1 % proliferation respectively: Figure 3).

**Figure 3.**
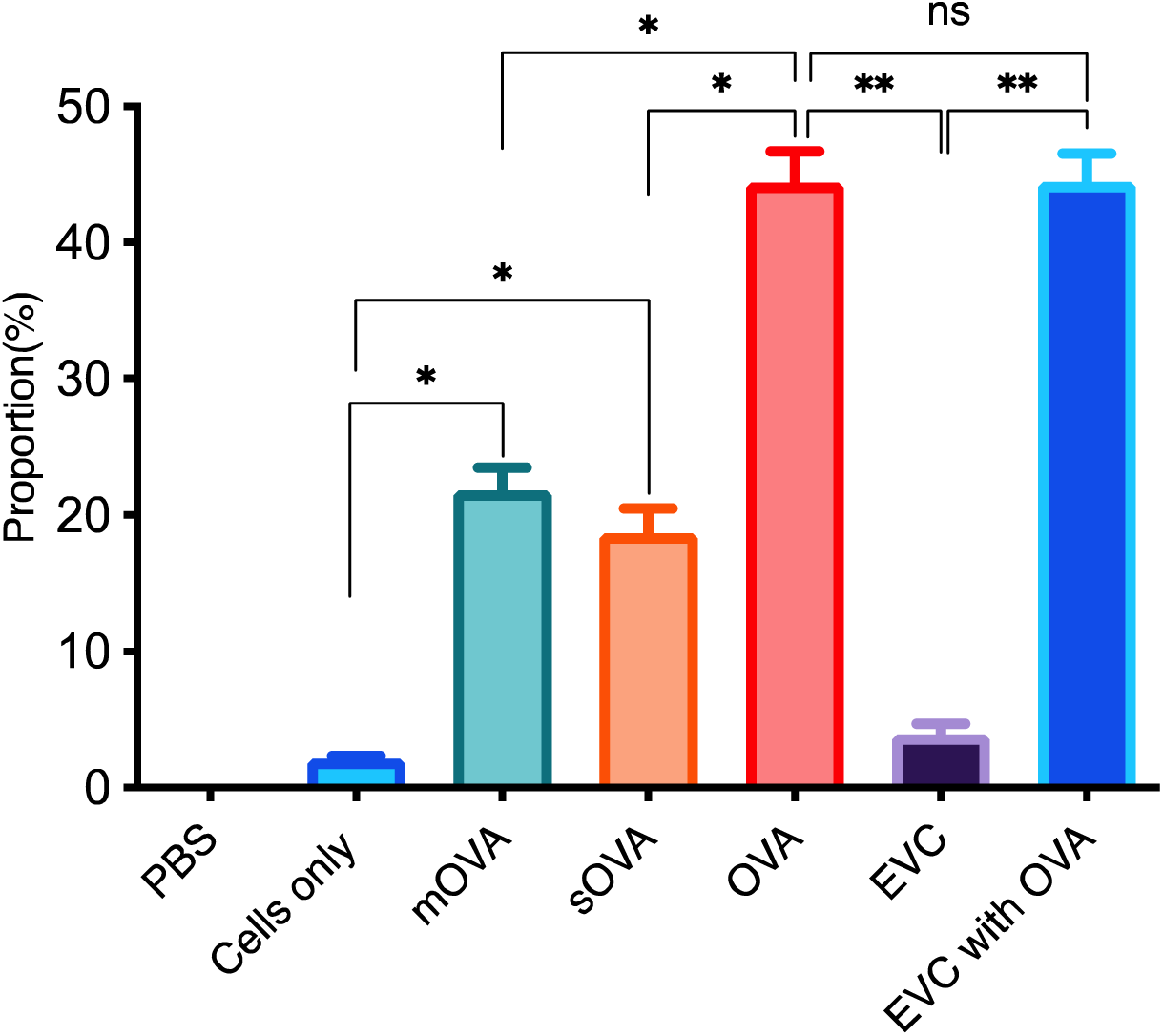
The proliferation of OT-II T cells following stimulating with OVA-expressing *L. major* parasites *in vitro*. A single cell suspension was prepared from spleen, maxillary, axillary, and inguinal lymph nodes of OT-II mice for an antigenic stimulation of the whole fraction or non-purified cells were labelled with cell division tracking dye, CFSE (0.5 μΜ) to a total of 2×10^5^ cells per well in 96 well plate, in a total of 8 wells for each parasite strains and after 4 days of incubation, the pool of cells were analysed for proliferation using PE-anti-mouse CD4 marker (Bio Legend). Proliferation was analysed using flow cytometry relaying on the CFSE dye being diluted following each cell division. EVC stands for empty vector control, mOVA stands for membrane-associated OVA, sOVA stands for soluble OVA. *, p<0.05; **, p<0.01; ns, not significant.

### 3.4. Expression of Th cytokines following activation with OVA-expressing *L. major*

The cytokine profiles of OT-II CD4^+^ T cells after exposure to OVA-expressing *L. major* was analysed in the culture supernatants of these cells. A total of seven cytokines representing the three T helper cell lineages namely, Th1, Th2 and Th17 were analysed. The result revealed that a significant amount of IFN-γ, TNF, IL-17, and IL-6 was detected in cells stimulated with mOVA and sOVA-expressing parasites compared to those cells stimulated with an equal number of parasites transfected with the empty vector (as a control) and cells left unstimulated (Figure 4). The expression of IL-2, and IL-10, was significantly higher in cells treated by OVA than mOVA and sOVA expressing parasites. In addition, the comparison in the expression level of all the cytokines between cells stimulated to mOVA and sOVA showed not statistically significant (Figure 4).

**Figure 4.**
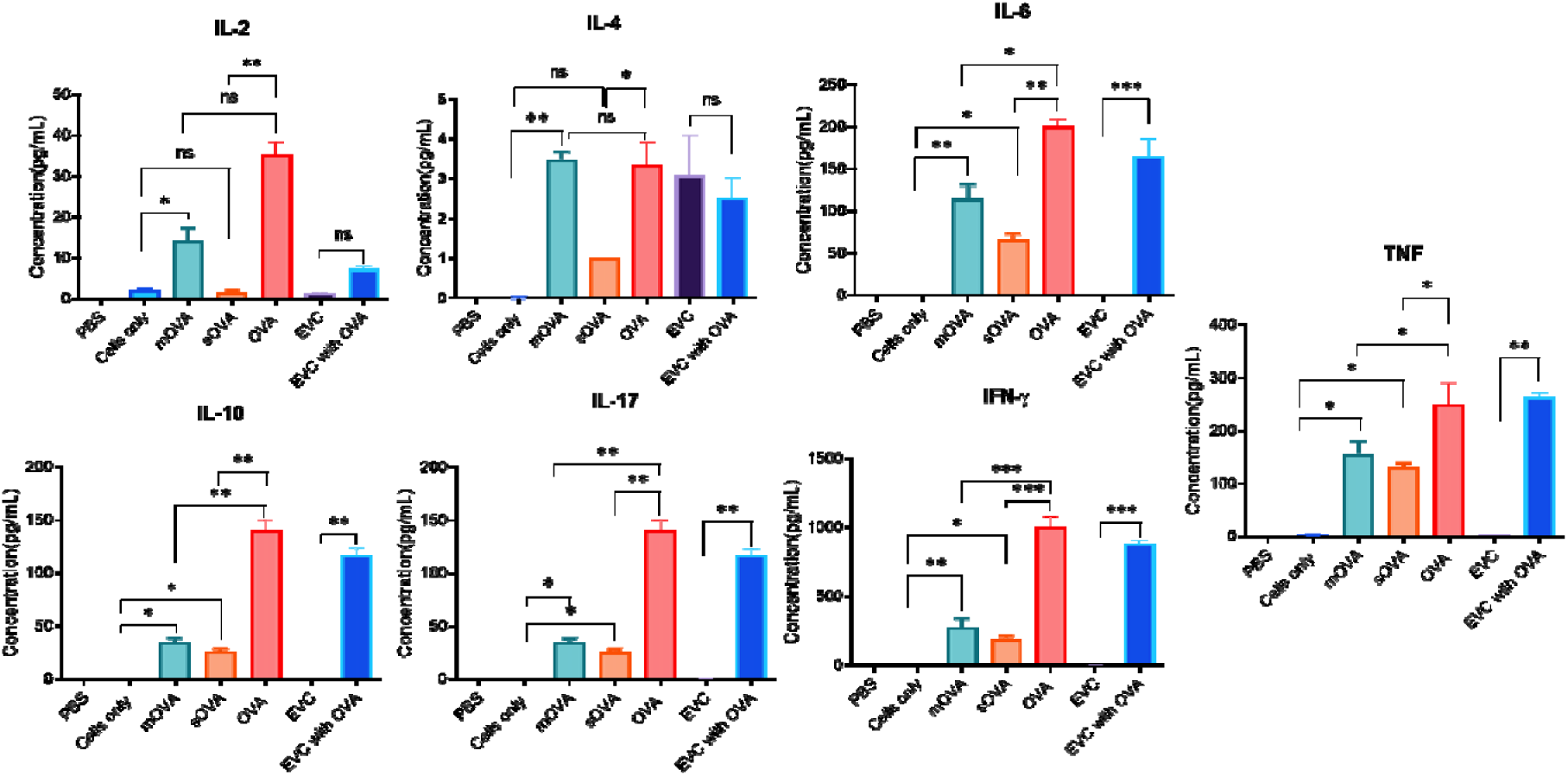
Detection of cytokines released by OT-II T cells stimulated with OVA-expressing parasites. A single cell suspension was prepared from spleen, maxillary, axillary, and inguinal lymph nodes of OT-II mice for an antigenic stimulation of the whole fraction or non-purified cells. A total of 2×10^5^ OT-II T cells per 96 well plates were cultured with each parasite strain and after 4 days of incubation, culture supernatants were taken to analyse the secretion of cytokines using cytometric bead array. EVC stands for empty vector control, mOVA stands for membrane-associated OVA, sOVA stands for soluble OVA *, p<0.05; **, p<0.01; ns, not significant.

### 3.5. OT-II T cells response to OVA expression *L. major*

The recognition of OVA-expressed by *L. major* promastigotes was analysed following transferring of 2×10^6^ CFSE-labelled OT-II CD4^+^ T cells into CD45.1 recipient or congenic mice. After 24 hours, mice were subcutaneously injected with 5×10^6^ mid-log phase transgenic parasites. Mice were injected with OVA peptide (10 μg) as a positive control. After one week, cells were extracted from the spleen and lymph nodes of the recipient mice and analysed for their ability to proliferate. The result shows a relatively higher percentage of proliferated cells recovered from mice injected with mOVA (3.53%) and sOVA (2.57%) compared to mice receiving only cells and mice injected with empty vector control (0.84% and 0.89% respectively; Figure 5). The proportion of proliferating cells in mice that received OVA peptide was 4.54% (Figure 5).

**Figure 5.**
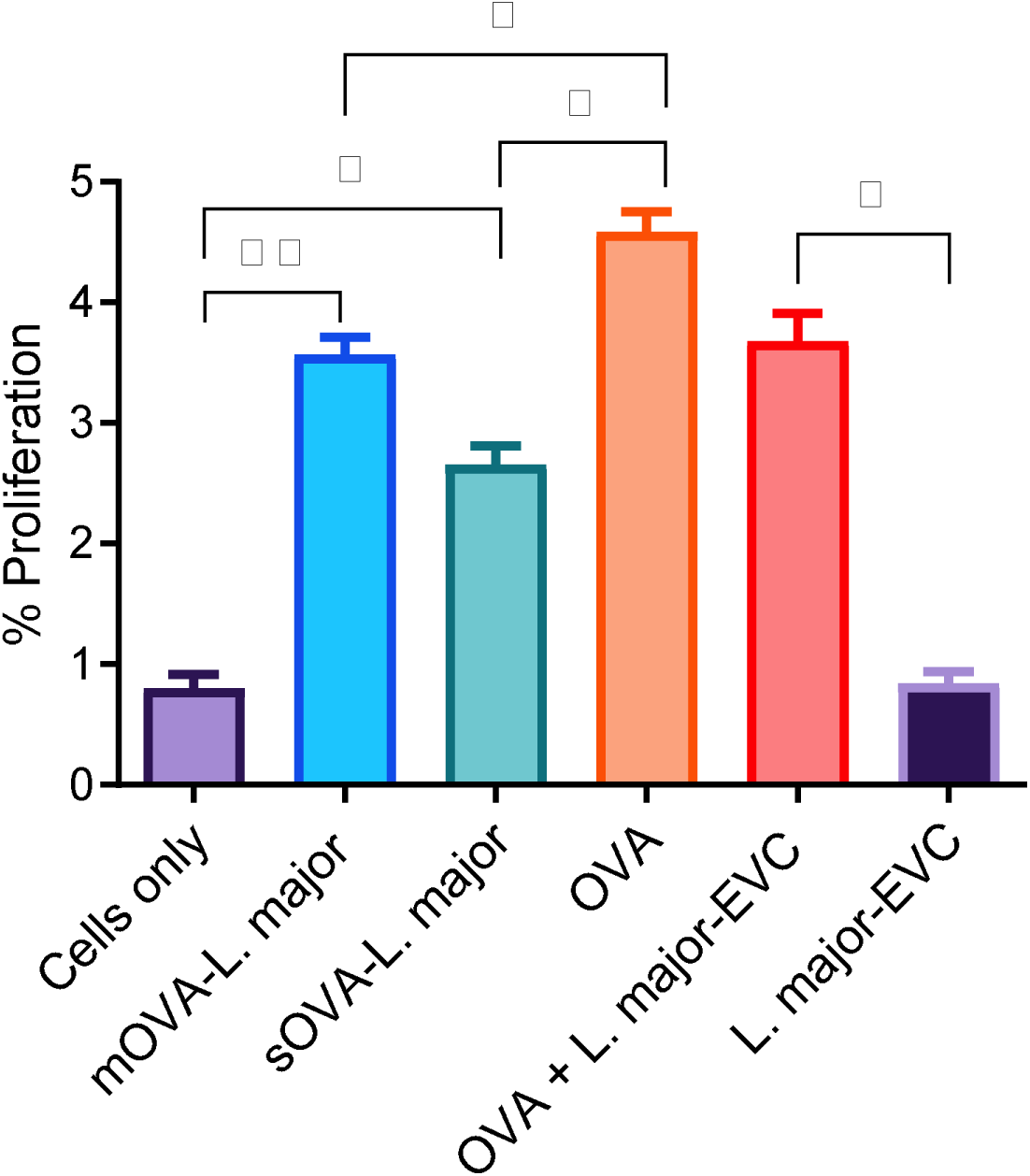
Proliferation rate of recovered OT-II T cells after parasite challenge. A single cell suspension was prepared from spleen, maxillary, axillary, and inguinal lymph nodes of OT-II mice for an antigenic stimulation of the whole fraction or non-purified cells. After injection of 2×10^6^ CFSE-labelled OT-II cells into congenic CD45.1-mice, 5×10^6^ OVA-expressing *Leishmania* promastigotes, and empty vector transformed promastigotes (EVC) were given s.c to each mouse. OVA (20μg/mL) and CFSE labelled cells without challenge were used as positive and negative control. The time between the first day of cell injection and challenge with the antigens was 24 hours. After the challenge, infection was maintained for a week before the mouse was culled for cell recovery and cells were extracted from the spleen and lymph nodes. After a week, recovered cells were directly analysed. EVC stands for empty vector control transfected *Leishmanial* parasite. *, p<0.05; **, p<0.01; ns, not significant.

### 3.6. *Ex vivo* response of OT-II T cells to OVA-expressing *L. major*

As described in the above section, approximately 2×10^6^ naive, unpolarized and CFSE-labelled OT-II T cells were intravenously injected into recipient mice and after 24 hours, each group of mice was challenged by s.c. injections with the following antigens: OVA protein (10 μg) as a positive control, 5×10^6^ parasites mixed with the same concentration of OVA, 5×10^6^ OVA-expressing log phase promastigotes (mOVA or sOVA), the same quantity of empty vector transformed parasites (EVC), and mouse injected with naïve or unpolarized cells left unchallenged to serve as a negative control. After a week, cells were recovered from spleen and lymph nodes of the recipient mice and further stimulated *ex vivo* with the same antigen used for the challenge and incubated for four days. After recovery from the recipient mice, injected OT-II T cells were identified from the recipient cells using anti-mouse CD45.2 marker (the recipient mice are CD45.1). Then cells were analysed for the proportion of cells proliferating and the results, as shown in Figure 6, demonstrate that a significant number of recovered cells proliferated when stimulated with OVA-expression *Leishmania* parasites. Only a marginal number of cells proliferated when those cells were recovered from mice challenged with the EVC-transformed parasite alone (0.5%) and remained unchallenged (i.e. no antigen-controls, 0.5%). In contrast, OT-II T cells recovered from mice challenged with parasite expressing mOVA, parasite expressing sOVA, OVA protein, OVA protein mixed with the parasite, had a proliferation rate of 3.7%, 2.8%, 5.0%, 4.6% respectively. Using a t-test, the proliferation rate of recovered cells stimulated *ex vivo* with OVA-expressing *L. major* (parasite expressing both forms of OVA) showed a significantly higher proliferation rate (p<0.05) compared to those recovered cells stimulated cells only or empty vector transformed parasites (Figure 6). No statistical significance was observed among recovered cells *ex vivo,* stimulated with mOVA-expressing parasite and sOVA-expressing parasite (p<0.05, Figure 6).

**Figure 6.**
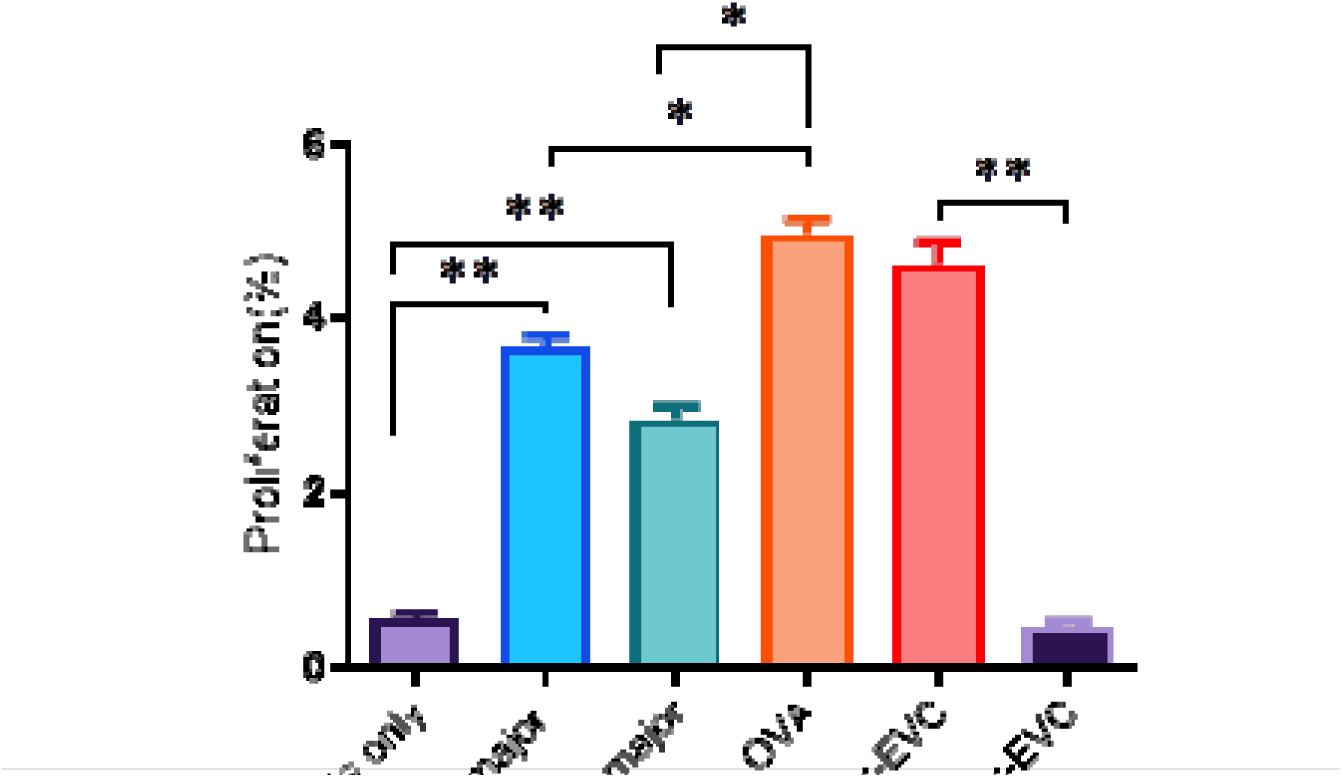
Proliferation rate of recovered cells following *ex vivo* stimulation. A single cell suspension was prepared from spleen, maxillary, axillary, and inguinal lymph nodes of OT-II mice for an antigenic stimulation of the whole fraction or non-purified cells. After injection of 2×10^6^ CFSE labelled OT-II cells into the congenic CD45.1-mice, 5×10^6^ OVA-expressing *Leishmania* promastigotes and empty vector transformed promastigotes (EVC) were given subcutaneous (s.c) to each mouse. OVA (10μg) and CFSE labelled cells without challenge were used as positive and negative control. The time between the first day of cell injection and challenge with the antigens was 24 hours. After a challenge, infection was maintained for a week before mouse culled for cell recovery. After a week, cells were extracted from the spleen and lymph nodes and were stimulated *ex vivo* with all antigens used for challenge separately for 12 hours. EVC stands for empty vector control transformed *Leishmanial* parasite. *, p<0.05; **, p<0.01; ns, not significant.

### 3.7. Cytokine production of recovered cells following stimulation with OVA-expressing *L. major*

To confirm the *in vivo* and *ex vivo* activation of OT-II cells recovered from congenic mice, we analysed the cytokine secretion profile of OT-II cells recovered from congenic mice and *ex vivo* stimulated. As shown in Figure 7, seven cytokines, namely IL-2, IL-4, IL-6, IL-10, IL-17, TNF, and IFN-γ, were produced by cells stimulated by the mOVA and sOVA. As expected, OT-II T cells recovered from mice challenged or immunized with OVA protein followed by *ex vivo* stimulation with the same protein also showed a significant level of cytokine expression. Whereas cells which were not challenged with any of the antigens, and those, challenged with empty vector transfected *Leishmanial* parasites (EVC) showed background-level expression of cytokines. For cytokines the concentration in supernatant of cells stimulated with OVA-expressing parasites (mOVA and sOVA) were significantly higher (P<0.05) than the cytokine expressed by cells exposed to no stimulation (cells only) or empty vector transfected parasites (EVC)(Figure 7).

**Figure 7.**
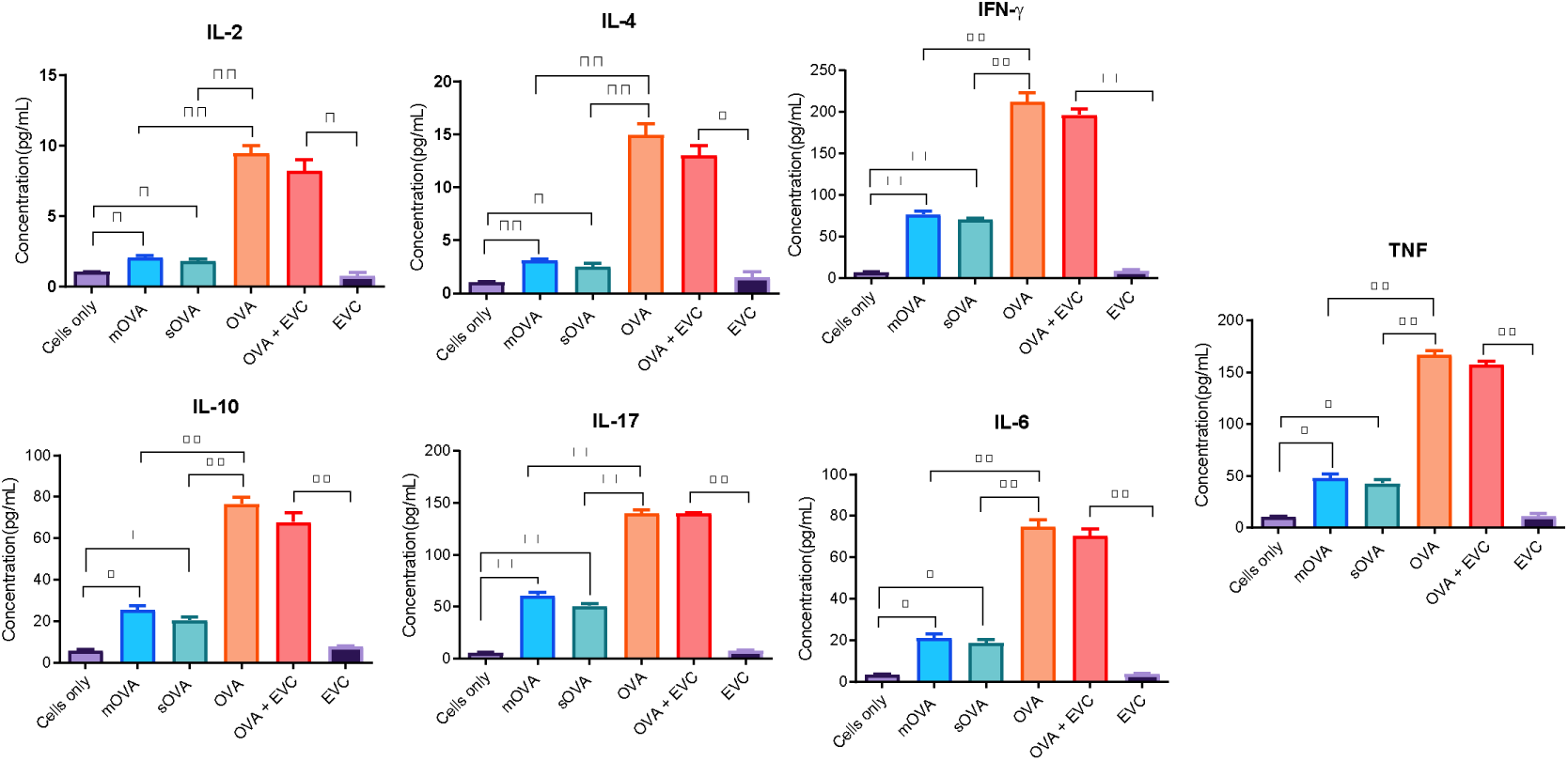
Cytokine expression of recovered cells following *ex vivo* stimulation. A single cell suspension was prepared from spleen, maxillary, axillary, and inguinal lymph nodes of OT-II mice for an antigenic stimulation of the whole fraction or non-purified cells. After injection of 2×10^6^ CFSE labelled OT-II cells into the congenic CD45.1-mice, 5×10^6^ OVA-expressing *Leishmania* promastigotes and EVC-transformed promastigotes (EVC) were given s.c to each mouse. OVA (10 μg) and CFSE labelled cells without challenge were used as a negative control. The time between the first day of cell injection and challenge with the antigens was 24 hours. After a challenge, infection was maintained for a week before the mice were culled for cell recovery. After a week, recovered cells were stimulated *ex vivo* with all antigens used for challenge. Cells were further cultured *in vitro* for four days, and culture supernatants were analysed for the expression of cytokines using cytokine bead array and flow cytometry. The concentration was expressed in pg/mL. EVC stands for empty vector control transfected *Leishmania* parasite. *, p<0.05; **, p<0.01; ns, not significant.

### 3.8. Stimulation and proliferation rate of recovered antigen-experienced cells

In the previous section, naïve OT-II cells were stimulated with various forms of OVA, but these cells were not polarised prior to adoptive transfer. To understand if *L. major* affects the Th phenotype of the OT-II cells and hence the degree of resilience of these cells, it was essential to polarise the OT-II cells prior to adoptive transfer and demonstrate that these cells can be stimulated by OVA-expressing *L. major*. Therefore, naïve OT-II T cells were polarized *in vitro* into different Th1, Th2, and Th17 cells using polarizing cytokines, antibodies to cytokines and OVA peptide and differentiated into memory cells using IL-7. These cells were injected intravenously into recipient mice and after 24 hours, each group of the recipients were challenged subcutaneously with different antigens: OVA protein (20μg/mL) as a positive control, 5×10^6^ empty vector transformed promastigotes mixed with the same concentration of OVA (20μg/mL), 5×10^6^ OVA-expressing log-phase promastigotes, same quantity of empty vector control transfected *Leishmania* parasites (EVC). In addition, mice injected with each phenotype of polarized cells were left unchallenged to serve as a negative control. After a week, memory OT-II T cells were recovered from spleen and lymph nodes of the congenic mouse and further stimulated *ex vivo* with the same antigen used for a challenge and incubated for four days. In all cases (i.e., Th1, Th2, and Th17) polarised OT-II T cells proliferated when stimulated with OVA (10 μg), mOVA-, and sOVA-expressing parasites (Figure 8). The result showed that following *ex vivo* stimulation, OVA protein induced a significantly higher amount of proliferation compared to both types of OVA-expressing parasites (Figure 8). Similarly, recovered cells of each phenotype showed a higher percentage of proliferation when stimulated with OVA-expressing *Leishmania* parasites compared to empty vector transfected *Leishmania* parasite and when cells were left without antigen stimulation (Figure 8). A marginal number of cells showed proliferation when recovered from mouse challenged with the EVC transfected parasites alone and those that remained unchallenged with any antigen.

**Figure 8.**
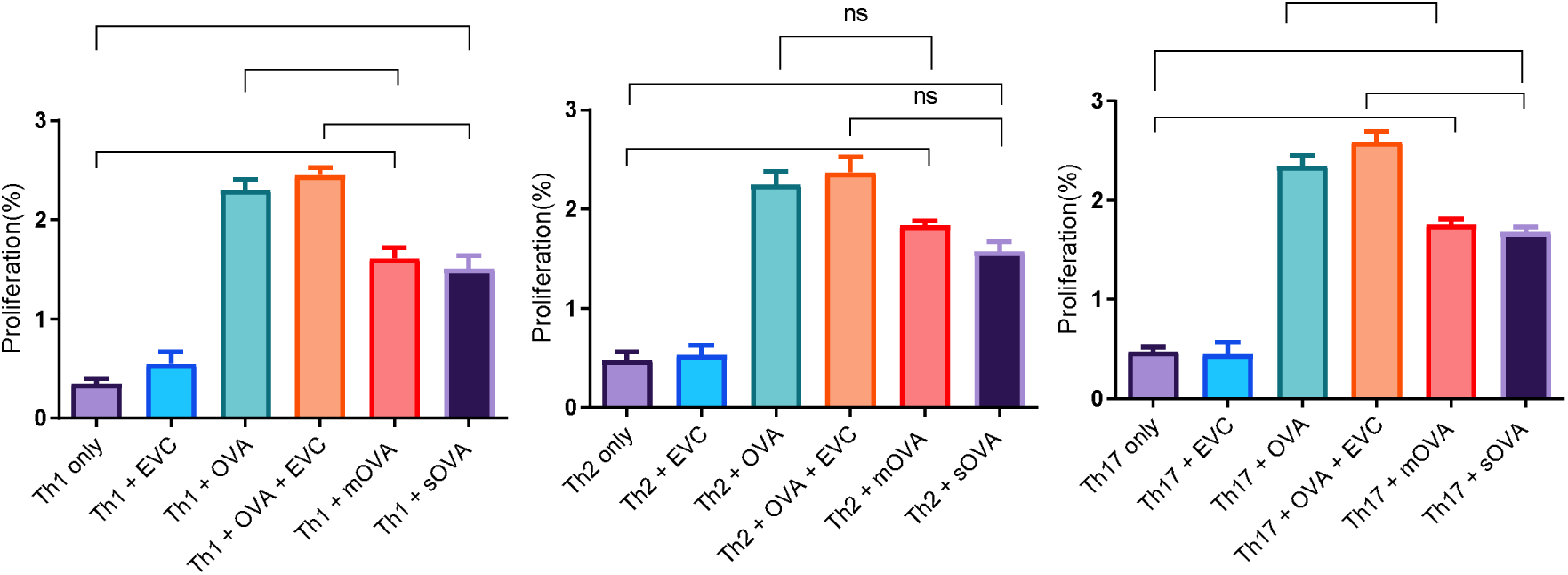
Proportion of proliferating polarised OT-II cells following *ex vivo* stimulation. A single cell suspension was prepared from spleen, maxillary, axillary, and inguinal lymph nodes of OT-II mice for an antigenic stimulation of the whole fraction or non-purified cells. After injection of 2×10^6^ polarised OT-II cells into the congenic CD45.1-mice, 5×10^6^ OVA-expressing *Leishmania* promastigotes, and EVC-transformed promastigotes (EVC) were given s.c to each mouse. OVA (10μg) and cells without challenge were used as a positive and negative controls, respectively. The time between the first day of cell injection and challenge with the antigens was 24 hours. After the *in vivo*-challenge, infection was maintained for one week before the OT-II cells were recovered for *ex vivo*-restimulating for 12 hours. The proliferation rate was estimated using flow cytometry by dividing the total proliferated and anti-CD4 marker positive T cells to total CD4^+^ T cells recovered. EVC stands for empty vector control transformed *Leishmanial* parasite. *, p<0.05; **, p<0.01; ns, not significant.

### 3.9. Cytokine production by recovered Th polarised cells following stimulation with OVA-expressing *L. major*

After *in vitro* polarization of naïve OT-II T cells into Th1, Th2, and Th17 using polarizing cytokines and OVA peptide (10 μg/mL), memory cells were produced using IL-7. Then approximately 2×10^6^ naive, Th1, Th2, and Th17 polarized OT-II T cells were injected intravenously into recipient mice and after 24 hours, each group was challenged by subcutaneous injections with the following antigens: OVA protein (10 μg) as a positive control, 5×10^6^ parasites mixed with the same concentration of OVA, 5×10^6^ OVA-expressing log-phase promastigotes, same quantity of empty vector transformed *Leishmania* parasites (EVC). In addition, some mice were injected with each phenotype of polarized cells and left unchallenged to serve as a negative control. After one week, antigen-experienced cells were recovered from spleen and lymph nodes of the congenic mice and further stimulated *ex vivo* with the same antigen used for a challenge and incubated for four days. Then cells were analysed for the production of cytokines as shown in Figure 9. Different cytokines were produced at various concentrations. The result showed that for the negative controls (i.e., cells only and EVC) there was either a non-detectable amount of cytokine or marginal amounts. Naïve cells stimulated with OVA or EVC + OVA, produced each of the cytokine tested. Naïve cells stimulated with mOVA or sOVA produced less of each cytokine compared to when they were stimulated with OVA. Interestingly, the cytokine profile of the differentiated OT-II cells whether Th1, Th2 or Th17 retained their cytokine secretion profile when stimulated with OVA or EVC + OVA (Figure 9). Indeed, Th1 cells continue to produce IFN-γ, irrespective of the stimuli (Figure 9). Th2 cells continue to produce predominantly Th2 cytokines such as IL-2, IL-4, IL-6, and IL-10 irrespective of the stimuli (Figure 9). Similarly, Th17 cells produce IL-17 at a higher level than any of the other differentiated OT-II cells (Figure 9). Conversely, Th2 and Th17 cells make very little IFN-γ, while Th1 cells make very little Th2 or Th17 cytokines. Beside IL-17, Th17 cells do not make any other cytokines to any great extent. Thus, irrespective of the stimuli provided the differentiated cells continue to make the cytokines that they were previously differentiated to make.

**Figure 9.**
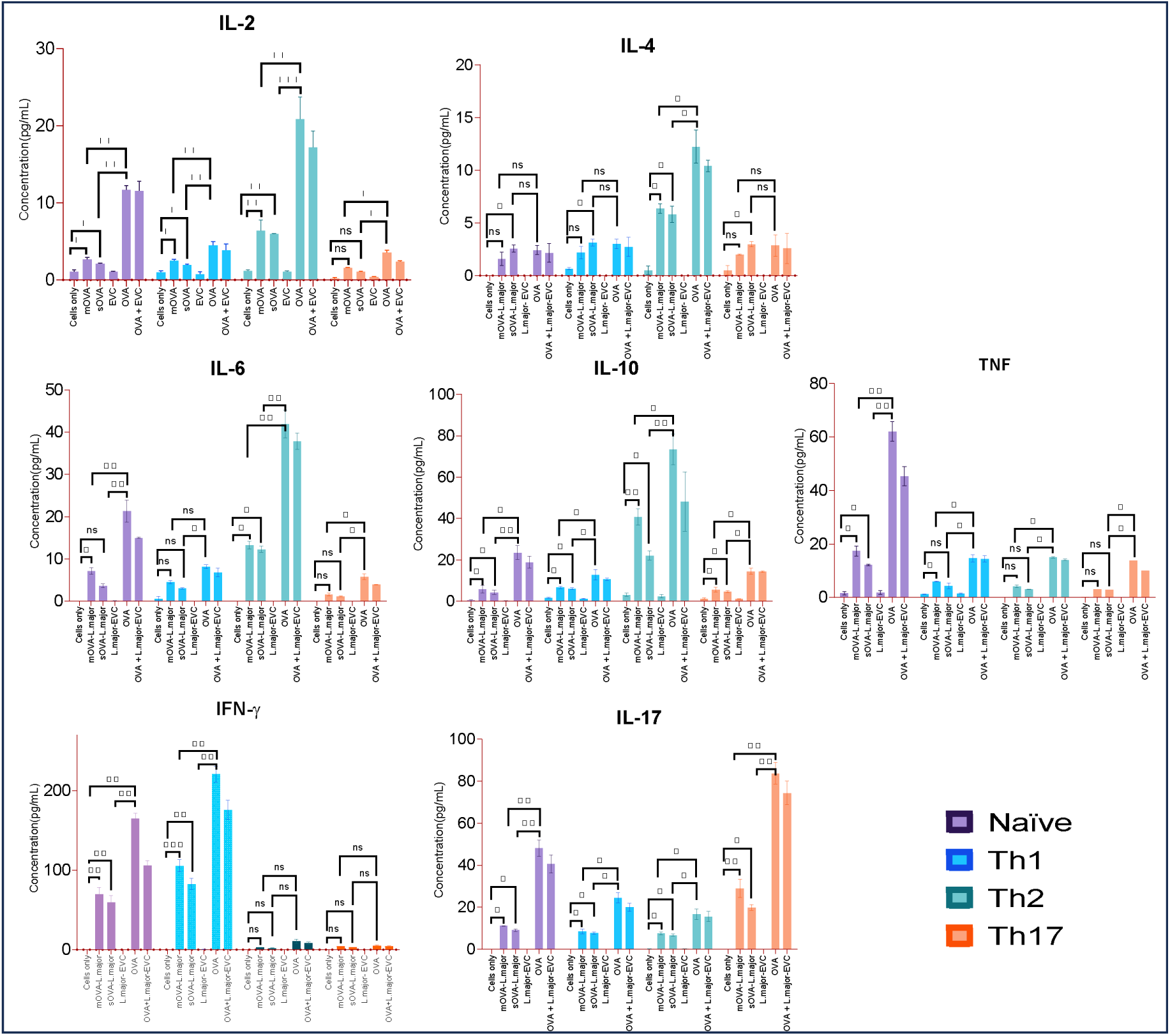
Detection of cytokine signatures from polarised OT-II T cells following antigen-specific *ex vivo* stimulation. A single cell suspension was prepared from spleen, maxillary, axillary, and inguinal lymph nodes of OT-II mice for an antigenic stimulation of the whole fraction or non-purified cells. After injection of 2×10^6^ Th phenotype differentiated OT-II cells into the congenic CD45.1-mice, 5×10^6^ OVA-expressing *Leishmania* promastigotes, and empty vector transformed promastigotes (EVC) were given s.c. to each mouse. OVA (10 μg) and cells without challenge were used as a positive and negative control respectively. The time between the first day of cell injection and challenge with the antigens was 24 hours. After a challenge, infection was maintained for a week before mouse culled for cell recovery. After a week, recovered cells were stimulated *ex vivo* for 12 hours with all antigens used for challenge separately. Then cells were further cultured *in vitro* for four days, and culture supernatants were analysed for the expression of cytokines using flow cytometry.

### 3.10. Measuring immune memory resilience during *L. major* exposure

In this study, the immune memory resilience of antigen-experienced cells (i.e., the ability to withstand manipulation by pathogens) was measured *in vivo* by analysing the cytokines in response to challenge to *L. major.* The result showed all antigen-experienced cells produce the same cytokine signatures as analysed using flow cytometry. As illustrated in Figure 10, following injection of congenic mice with naïve unpolarized cells, Th1, Th2 and Th17 polarized cells, each group of mice were challenged either with OVA (10 μg), mOVA-expressing *L. major* and finally recovered unpolarized (naïve) and antigen-experienced cells were further cultured *in vitro* by stimulating with OVA (10 μg/mL). The memory cells were analysed to measure their cytokine signatures and the result showed that mice injected with naïve cells produce a lower level of cytokines (less than 10 pg/ml); whereas, mice injected with Th1 phenotypes followed by challenge with OVA and mOVA-expressing *L. major* produce their cytokine signatures with concentrations of IFN-γ (65.2 pg/ml; 50.1 pg/ml) and TNF (15.8 pg/ml; 13.2 pg/ml) respectively. In addition, mice injected with Th2 phenotypes followed by challenge with OVA and mOVA-expressing *L. major* produce IL-10 (32.1 pg/ml; 22.4 pg/ml) and IL-6 (21.2 pg/ml; 15.6 pg/ml) respectively. Similarly, mice injected with Th17 phenotypes followed by challenge with OVA and mOVA-expressing *L. major* produce IL-17 (28.2 pg/ml; 19.6 pg/ml) and TNF (11.8 pg/ml; 9.4 pg/ml) respectively. The statistical association between the cytokines produced by the primed cells comparing to the non-primed or naïve cells following exposure to antigen stimulation showed Th1 cytokine except TNF, Th2 and Th17 cytokines expressed significantly higher level comparing to unprimed cells following antigenic stimulation with mOVA expression parasites. In comparison, naïve OT-II T cells produced a minimum number of cytokines following exposing to OVA-expressing parasites and OVA protein (Figure 10).

**Figure 10.**
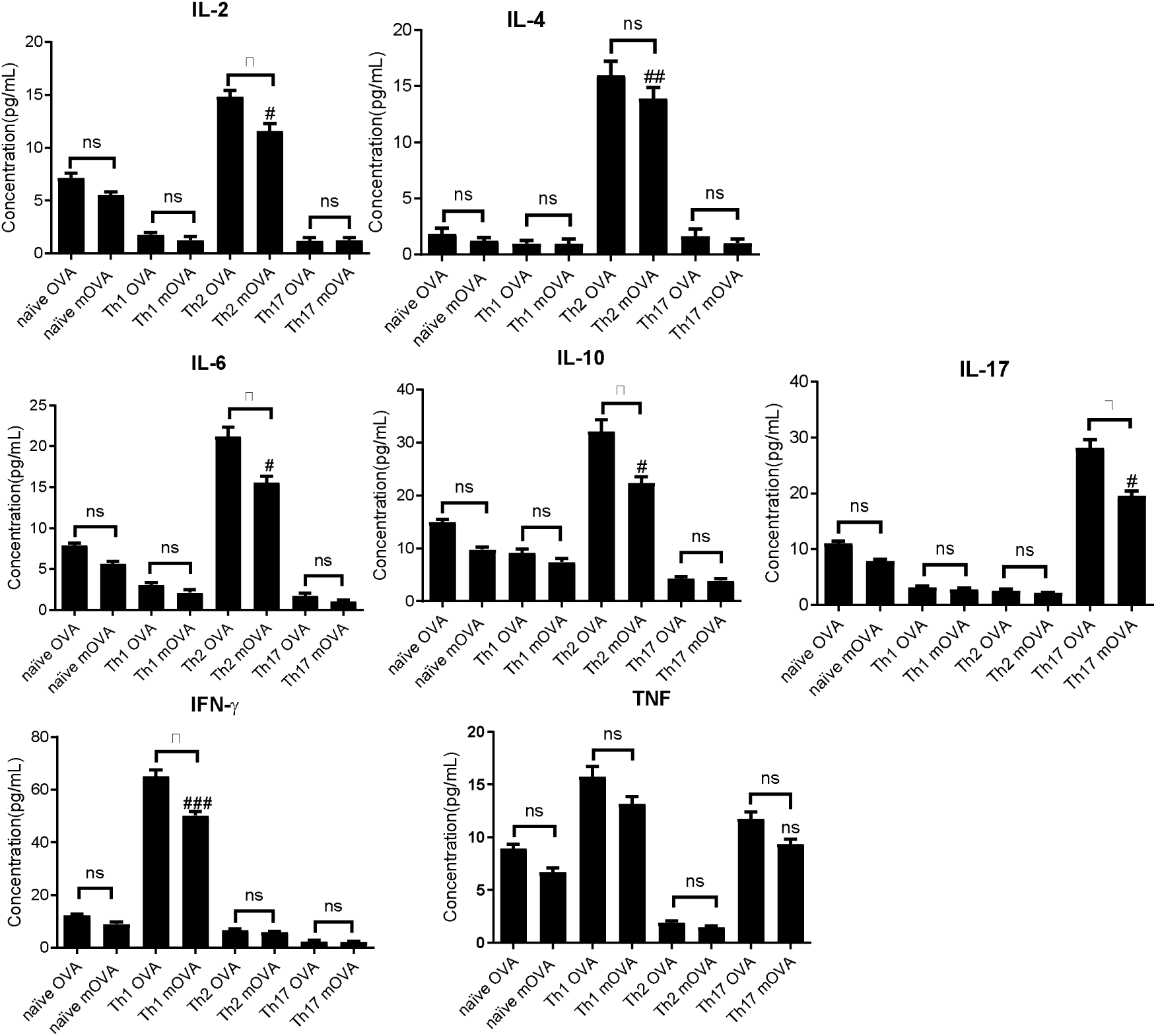
Cytokine signatures of antigen-experienced cells which underwent *in vivo* challenge with OVA transfected parasites or cells only control, following *ex vivo* stimulation with OVA protein. A single cell suspension was prepared from spleen, maxillary, axillary, and inguinal lymph nodes of OT-II mice for an antigenic stimulation of the whole fraction or non-purified cells. Then, naïve OT-II T cells were polarized into Th1/ Th2/ Th17 phenotypes. Then, a total of 2×10^6^ cells were transferred into recipient congenic mice (CD45.1, n=6, iv) and after 24 hours, each mouse was challenged with 5 × 10^6^ log-phase promastigotes (mOVA-expressing *L. major*) and OVA protein (20 μg OVA). Control mice received naïve undifferentiated OT-II T cells. After seven days, recipient mice were terminated, and memory antigen-experienced cells were processed and restimulated with 2μg OVA. Finally, cytokine signatures were analysed in the culture supernatant.

## 4. Discussion

The efficacy of experimental *Leishmania* vaccines is significantly influenced by innate immune cells. For example, neutrophils play a critical role in vaccine immunity due to their antigen presenting capacity to naïve T cells. Skin resident CD4+ memory T cells recruit Inflammatory monocytes(iMO) and contribute to controlling *L. major* infection by producing inducible nitric oxide synthase and nitric oxide. During vaccine immunity, the function of macrophages in enhancing a Th1 or Th2 response has been reported in *L. donovani* infections. In addition, the role of DCs as a source of IL-12 & its role in the pathogenesis & inducing an efficacious vaccination response in *Leishmania* is also reported (31). Immunization using centrin deficient *L. mexicana* parasites (*LmexCen ^−/−)^*, following generation of CD4 + T memory cells showed a protective immunity can be maintained. These genetically manipulated parasites are potentially used for vaccination and can provide protection against *L. mexicana* infection (32). Changes in MHC-II/peptide affinity, antigen presentation, protease-susceptible sites, and intracellular expression of pathogenic proteins during *Leishmania* infection can affect the reactivation of targeted antigen-specific T cells produced during priming, as well as dominant epitope selection. When *Leishmania*-infected macrophages provide antigens to memory T cells that are specific to that antigen, the T cells functional phenotype may shift, resulting in either inactivity or death. The peptides produced during infection may differ from and cross-react with the priming peptides, even though cells may be stimulated. The otherwise active T lymphocytes that are specific to antigens may be suppressed by such modified peptide ligands (33). In this study, we hypothesized that a long-lasting protective immunity in a host can be induced through the development of immune memory T cells and by the ability of these cells to induce appropriate secondary response to specific pathogenic infections. This can be achieved through the generation of a resilient immune memory that can withstand the manipulation process by the parasite. Therefore, we generated *L. major* parasites, which express two forms of OVA namely, membrane-associated OVA (mOVA) and soluble OVA (sOVA). We showed that expression of both forms of *L. major*-expressed OVA can stimulate OT-II cells *in vitro* and *in vivo*. These results were unexpected because previous studies have shown that while mOVA could stimulate OT-II cells soluble (or cytosolic) sOVA could only at a higher dose (34). Indeed, Prickett et al.2006(34), used *L. major* hydrophilic acylated surface protein B for the generation of OVA transgenic parasite and expressing the antigen either in the cytosol or membrane associated. Only the membrane-associated OVA was recognized and leads to the activation of CD4^+^ T cells at low parasite numbers but both mOVA and sOVA-producing parasites could stimulate immune recognition at higher concentrations. Such studies support the importance of antigen localization during the recognition by T cells (35). However, in our experiments we used higher parasite doses, which could account for the observed stimulation of OT-II cells with sOVA at a similar level to mOVA. In addition, Prickett et al used DO11.10 TCR Tg mice (34), while we used OT-II TCR transgenic animals and the amount of accessible OVA required to stimulate these two different T cells might be different. Our results are compatible with these of Kaye et al. 1993(36), suggesting that *L. major* expressing OVA and β-gal in the phagosome as soluble antigen, being efficient at inducing CD4^+^ T cell activation. Nevertheless, in later experiments when number of animals were limiting, and we had to choose only one type of parasite, we chose mOVA.

Although the level of cytokines produced was generally much smaller than in the positive controls containing OVA protein, every cytokine measured was produced following stimulation with both sOVA and mOVA-expressing parasites. The relatively low level of cytokine produced was not due to an inhibitory or toxic effect of the parasites on the OT-II cells as control parasites supplemented with soluble OVA protein secreted cytokines at a similar rate to the OVA protein-stimulated OT-II cells. Thus, it is likely that the relatively low level of cytokine secretion is due to the relatively low amount of OVA-protein produced by the parasite and/or the way that the parasite-produced OVA, is processed within the APCs. Indeed, it has been suggested that the antigen processing of *L. major* produced OVA might be different to soluble OVA protein, as phagolysosome-targeted OVA expressed in *L. major* was able to be presented by macrophages for a period of 24h, unlike these same macrophages pulsed with soluble OVA (36). Therefore, if indeed OVA produced by the parasite can be presented for a prolonged period of time, it is likely that the concentration of peptides associated with the MHC II molecules on the surface of these cells at any given time might be lower than in the case of soluble OVA present in the surrounding environment.

We also demonstrated that OT-II cells were readily polarised *in vitro*, and these polarised OT-II cells behaved as expected and could be restimulated, not only by OVA but importantly also by *L. major*-expressing both mOVA and sOVA. This opened the door to *in vivo* experiments using OT-II cells transferred to recipient mice. This is important as intracellular parasites such as *Leishmania*, living inside the harsh environment of phagocytes have developed strategies and physiological adaptations to escape the immune system of the host. Some of these mechanisms involve the manipulation of the host’s immune response to achieve, for example, inhibition of the signalling pathways of the host’s macrophage leading to parasite survival (37). Different *in vitro* studies confirmed that IL-10, one of the anti-inflammatory cytokines, is produced by macrophages parasitized with *Leishmania* through interaction with Fcγ receptor (38). The production of IL-10 leads into the suppression of the microbicidal nature, which involves nitric oxide, production of cytokines such as TNF, IL-1, and IL-12. The expression of costimulatory molecules such as B7-1/2 has also been involved in the process of inhibiting the microbicidal activities of macrophages (39). The clinical implication of IL-10 expression has been studied *in vivo* and the results indicated that transgenic mice expressing IL-10 are failing to control the parasite (40). However, the finding from our study demonstrated that in the presence of Th polarized cells, the production of IL-10 is not associated with progression of *Leishmania* infection at a higher rate as the parasite load (Figure S1) was shown to be significantly reduced in mice injected with either Th1, Th2, or Th17 cells followed by a challenge with OVA expressing parasites and the parasite control. This was unexpected as the Th2 cells produced significantly more IL-10 compared to the Th1 and Th17 cells and hence we might have expected, based on previous studies (41, that the presence of IL-10 would have promoted parasite development. At this stage it is unclear why this might be the case.

The efficacy of experimental *Leishmania* vaccines is significantly influenced by innate immune cells. For example, neutrophils due to their antigen presenting capacity to naïve T cells, they play a critical role in vaccine immunity. Skin resident CD4+ memory T cells recruit Inflammatory monocytes(iMO) and contribute to controlling *L. major* infection by producing inducible nitric oxide synthase and nitric oxide. The role of DCs as a source of IL-12 and its role in the pathogenesis and potentiating an efficacious vaccine response in *Leishmania* is also reported (31). Several experimental vaccines against *Leishmania* have been generated to halt the skin and visceral forms of the disease, which are contracted through the bite of an infected sand fly. Following needle challenge, immunization with autoclaved L. major (ALM) given with the Th1 adjuvant TLR9 agonist CpG oligodeoxynucleotides significantly protects against infection in the resistant C57BL/6 mouse. In resistant C57BL/6 mice, an immunity produced by KSAC or L110f immunization with GLA-SE after challenge with L. major by needle or infected sand fly bite was investigated. Since sand fly bite is expected to induce neutrophil recruitment, the influence of KSAC vaccination on the strength and Th1/Th2 nature of subsequent immune responses to L. major transmitted by sand fly bite in these two mouse strains is likely to be in addition to any influence of neutrophils on the expression of this immunity (42). Following a challenge of *in vitro* polarized memory T cells by OVA-expressing *L. major* parasites *in vivo*, the degree of immune memory resilience was measured. Naïve OT-II T cells were polarized into Th1, Th2 and Th17 *in vitro* and these cells’ phenotypes were confirmed by analysing their cytokine signatures. These cells when adoptively transferred to recipient mice, and they retained their cytokine secreting profiles when restimulated *in vivo* and analysed *in vitro*. This is a key finding as it suggests that *in vitro* polarised OT-II cells can maintain their Th profile *in vitro* when subjected to *L. major* expressing OVA. This is extremely important for vaccine development as it suggests that if it were possible to vaccinate in such a way as to generate immune memory cells similar to these produced *in vitro*, one should be able to protect from infection. Adoptively transfer of in vitro differentiated Th1 cells specific for *Leishmania*-specific CD44^+^PEPCK tetramer into recipient mice showed immediate presence of circulating CD4^+^ T helper 1 effector cells (Th1_EFF_). This is critical to prevent *L*. *major* mediated immunomodulation (43). Experimental studies showed that understanding the link between immune memory and protection against a parasite is very important as it has the implications for efficacious vaccination. In addition, memory T cells are reported to be protective only if they generate circulating Teff cells before challenge (44). Two distinct populations of CD4(+) T cells, namely short-lived pathogen-dependent Teff and long-lived pathogen-independent Tcm define immunity to *L. major* (45).

Skin graft transplantation of *Leishmania* -specific CD4+ T memory cells on to naïve mice produce IFN-γ and remain resident in the skin. They recruit circulating T cells to the skin in a CXCR3-dependent manner and controls the parasite infection, and this draws a conclusion that optimal protective immunity to the parasite and effective vaccination depends on generating both circulating and skin-resident memory T cells (46). Tissue resident memory CD4^+^ T cells provide protection after parasite challenge and this requires recruitment and activation of inflammatory monocytes by producing reactive oxygen species and NO which plays a critical role in immunity to cutaneous leishmaniasis (47). After resolution of primary infection, short lived Teff and long lived Tcm cell, and skin resident memory T cells retained immunity to secondary infection. Although parasite infection leads to generation of protective immunity, vaccine efficacy depends on generating memory T cells which is sustained in the absence infection (48).

Indeed, induced protective immunity can be achieved when memory T cells can develop the ability to retain the production of their cytokine signatures on the face of the immunomodulatory pathogen. To this end, a resilient immune memory was detected *in vivo* after adoptive transfer of antigen-experienced cells into recipient mice and followed by challenge with *L. major*. Th1 polarized memory T cells showed the strongest proliferation and produced a significant amount of IFN-γ following challenge with OVA protein and OVA-expressing *L. major*. In addition, mice that received Th2 polarized memory T cells showed Th2 proliferation and produced a significant amount of IL-2, IL-4, IL-6 and IL-10 following challenge with OVA protein and OVA-expressing *L. major*. Similarly, mice which received Th17 polarized memory T cells showed OT-II cell proliferation and produced a significant amount of IL-17 following challenge with OVA protein and OVA-expressing *L. major*. due to several technical issues, they are using non-physiological number of parasites, T cells, and a non-physiological route of infection. One of the limitations of this study is the physiological number of parasites. In this study, due to technical issues related with transfection efficacy and optimizing the expression level of OVA protein by the parasites and induce OVA-mediated T cell activation, a relatively high number of parasites (5×10^6)^ was used compared to the number of parasites used in previously reported studies. To optimize the optimized the number to have an optimum level of protein expression to induce OVA-mediated T cell activation. Route of parasite infection is an important parameter to consider in different experimental models of *Leishmania* parasites. Although the most preferred route of parasite injection is intradermal to mimic natural infection and study immune response, evidence from previous study showed the subcutaneous route induces significantly more protective immunity against L. major, low parasite load, rapid lesion resolution, and lowers IL-production (49). However, we also recommend the intradermal route for the application of OVA transgenic parasite model for future immune modulation studies. In addion, the limitation of the study is related with non-optimal number of antigen experienced cells following the use of non-purified OT-II-T cells from spleen and lymph nodes. In this study, we have also used empty vector transfected parasites as a negative control but to explicitly see if there is an immune response induced by the parasite, including a wild type of group is recommended for future studies.

## 5. Conclusion

*In vitro* polarized Th1, Th2, and Th17 phenotypes remain resilient *in vivo* on the face of pathogen-mediated manipulation. The strongest immune memory response was observed in response to challenge with OVA protein or OVA-expressing *L. major* parasites as measured by the proliferation rate of recovered Th1 memory cells and the amount of IFN-γ production. This can reflect the fact that Th1 memory T cells have developed an intrinsic nature to strongly respond to OVA antigen in the face of immune manipulative pathogen because the recall response of Th1 memory cells to OVA_323-339_ was higher in terms of proliferative response and overall cytokine production as confirmed by the data in which mice received Th1 polarised memory T cells produced higher quantities of IFN-γ, but not other cytokines. In addition, findings regarding the effect of antigen localization on measuring the strength of T cell responses also showed that mice, which received Th1, Th2, and Th17 polarized cells and challenged with mOVA-expressing *L. major* revealed a relatively higher proliferative responses *in vitro* and *in vivo* compared to the mice that received the same group of antigen-experienced cells but were challenged with sOVA-expressing *L. major*. This is confirmed by measuring the proliferation of recovered memory T cells or by measuring the amount of cytokine they produced following restimulating with OVA protein.

## Acknowledgements

We are grateful to Melbourne International Research Scholarship for research training sponsorship. The authors also want to express their gratitude to Dr Fiona Sansom for her support in growing *Leishmania* parasites in vitro.

## Funding

No specific grant was received for this research from funding organizations in the public, private, or not-for-profit sectors.

## Availability of data and materials

The corresponding author can provide the datasets used and/or analyzed during the current work upon request.

## Authors’ contributions

MGT and MFN performed the experimental works, MGT analysed data, and drafted the paper, ALE and JYS reviewed and commented the draft paper. All authors read and approved the final version of the draft paper.

## Ethics approval

Ethical approval for animal studies was approved by the animal ethics and welfare committee of the University of Melbourne.

## Consent for publication

Not applicable

## Competing interests

The authors declare that they have no competing interests to disclose.

## Notes

### Competing Interest Statement

The authors have declared no competing interest.

